# Time-course analysis of the transcriptome of *Arabidopsis thaliana* leaves under high-concentration ammonium sulfate treatment

**DOI:** 10.1101/2024.05.02.591416

**Authors:** Hiroko Iwanaga, Yuki Arai, Maiko Nezuo, Akiko Doi, Takahito Takei, Masayuki Fujiwara, Takushi Hachiya, Takahiro Hamada

## Abstract

Nitrogen is essential for plant growth and is sourced primarily from nitrate and ammonium in the soil. Even though plants can take ammonium up for nutrition, it often results in toxic effects such as growth suppression and chlorosis. To elucidate the mechanism of ammonium toxicity, a time-course analysis of the transcriptome was performed on *A. thaliana* leaves treated with high concentrations of ammonium sulfate in the presence of sufficient nitrate. The expression of nitrate-inducible genes tended to be downregulated by the treatment. The expression of genes relating to abscisic acid, jasmonic acid (JA), salicylic acid (SA), and membrane trafficking was upregulated, whereas that of photosynthesis-, auxin-, and cytokinin-related genes involved in growth and development was downregulated. The induction of many osmotic stress-responsive genes suggests the involvement of osmotic stress in ammonium toxicity. Furthermore, the upregulation of nitric oxide (NO)-inducible genes and the simultaneous upregulation of genes involved in JA biosynthesis, glutathione metabolism, and SA response suggested the involvement of endogenous NO and protein S-nitrosylation in response to high concentrations of ammonium sulfate. This study provides a novel and comprehensive overview of transcriptional changes occurring in response to high ammonium sulfate concentrations and proposes possible mechanisms of ammonium toxicity that can be explored in future research.

## 1. Introduction

The availability of nitrogen (N) determines plant growth and development, as well as crop yield and quality. For most plants, nitrate and ammonium are the primary sources of N in the soil. In C_3_ plants, including major crops such as rice, wheat, and oilseed rape, nitrate reduction decreases with elevated atmospheric CO_2_ concentrations, whereas ammonium utilization remains unchanged [1]. Given that a decrease of as little as 1% in N use efficiency could increase the cost of crop cultivation worldwide by approximately USD 1 billion annually [2], ammonium fertilization is of great interest for future-oriented agriculture in a high-CO_2_ world. However, unlike nitrate, millimolar concentrations of ammonium often cause growth suppression and chlorosis [3,4]. Widely referred to as ammonium toxicity, many hypotheses have been proposed to explain the mechanism of this toxic phenomenon, including futile transmembrane ammonium cycling, deficiencies in inorganic cations and organic acids, impaired hormonal homeostasis, disordered pH regulation, defects in protein *N*-glycosylation, redox imbalance/oxidative stress, and the uncoupling of photophosphorylation [5,6]. In addition, recent efforts using *A. thaliana* and rice have identified some genetic loci that alter ammonium sensitivity [7–10].

Nevertheless, due to the involvement of ammonium toxicity in various physiological processes, a comprehensive understanding of the mechanism of toxicity remains elusive. To address this lack of understanding, we conducted a time-course analysis of transcriptome in *A. thaliana* leaves treated with high concentrations of ammonium. We selected ammonium sulfate as the ammonium source, given that it is one of the most common ammonium-containing fertilizers worldwide. Additionally, we focused on leaves because they are the primary photosynthetic organs in plants, playing a pivotal role in crop yield. Understanding the mechanisms of ammonium toxicity in leaves is therefore indispensable for the effective utilization of ammonium fertilizers in crop cultivation.

Accordingly, this study evaluated the effects of the combination of ammonium and sulfate on the leaf transcriptome. High concentrations of ammonium sulfate induce genome-wide transcriptomic reprogramming in genes related to photosynthesis, phytohormone biosynthesis and signaling, nitrate/N starvation response, nitric oxide (NO) response, osmotic stress response, sulfur assimilation, and membrane trafficking machinery. In addition, possible interactions among the responses and their relevance to ammonium toxicity are discussed.

## 2. Materials and Methods

### 2.1. Plant growth conditions

Six seeds of wild-type *Arabidopsis* (*Arabidopsis thaliana* Col-0) and a JA synthesis impaired mutant (*dde2-2*) [11] were surface-sterilized and sown on each piece of solidified medium, covered with a cellophane disk, in a 9 cm dish to minimize root wounding when transplanting between different media [12]. The medium contained approximately 30 mL half-strength Murashige and Skoog (1/2 MS) salts without N, supplemented with 20 mM KNO_3_, 5 mM (NH_4_)_2_SO_4_, 2 g L^−1^ MES, 1% (w/v) sucrose, and 0.25% (w/v) gellan gum (Fujifilm Wako, Osaka, Japan). The pH was adjusted to 5.8 with KOH. The sown seeds were incubated in the dark at 4°C overnight to enhance the breaking of dormancy. Plants were grown horizontally under a PPFD of 50–60 μmol m^−2^ s^−1^ (16/8-hour light/dark cycle) at 22°C.

### 2.2. High ammonium sulfate treatment

For the high ammonium sulfate treatment, three types of media with different concentrations of ammonium sulfate were prepared. The control medium contained the same composition as the cultivation medium (5 mM (NH_4_)_2_SO_4_), and the high ammonium sulfate media contained the same composition as the cultivation medium except for a higher ammonium sulfate concentration of 12.5 mM or 37.5 mM (NH_4_)_2_SO_4_. After 21 days of growth, the cultivated plants-on-cellophane were directly transferred onto either medium and then grown for another 24 h. Subsequently, the plants were retransferred onto the fresh control medium again to observe recovery from the treatment. Leaf samples were taken at 3, 6, and 24 h after the treatment, and at 24 h after plants were returned to the fresh control medium.

### 2.3. Sampling for ammonium concentration measurements and RNA-seq

For tissue ammonium measurement and RNA-seq, three independent plants in the same dish were pooled together to form each biological replicate. For each time point and each (NH_4_)_2_SO_4_ concentration, two replicates were collected from two independent dishes. For each replicate, the first true leaves of the three pooled plants were collected for tissue ammonium measurement, while the remaining true leaves from the same pooled three plants were collected for RNA extraction. The samples for ammonium measurement were weighed and transferred to 2 mL polypropylene tubes (Yasui Kikai, Osaka, Japan) containing zirconia beads (5 mm in diameter), and the samples for RNA extraction were transferred to 1.5 mL tubes (WATSON Bio Lab, Tokyo, Japan). The tubes were immediately frozen in liquid N_2_ and stored at −80°C until use.

### 2.4. Measurements of the ammonium concentration

Frozen samples were ground with a Multi-Beads Shocker (Yasui Kikai) using zirconia beads (5 mm diameter). The frozen powder was added to 10 volumes of 0.1 N HCl and mixed vigorously at room temperature for 10 min. The mixture was centrifuged at 20,400 × *g* at 4°C for 5 min. The supernatant was filtered through Ultrafree-MC Centrifugal Filter Units (No. UFC30GV00, Merck, Tokyo, Japan) at 12,000 × *g* and 4°C for 2 min. The ammonium concentration of the filtrate was determined using an automated amino acid analyzer (LaChrom Elite, Hitachi High-Tech Corp., Japan) in accordance with the manufacturer’s instructions.

### 2.5. RNA extraction and sequencing

Total RNA was isolated from frozen samples using the ISOSPIN Plant RNA kit (NIPPON GENE, Tokyo, Japan) in accordance with the manufacturer’s protocol. Approximately 3–12 µg (>120 ng/µL) of total RNA for each sample was sent to Novogene Japan K.K. for quality control, library preparation, and RNA sequencing. Paired-end sequencing with 150 bp reads for each end was performed using an Illumina NovaSeq 6,000 sequencer, ensuring a minimum sequencing depth of 4.9 G bases and 33 million reads per sample. The sequence data were deposited in the DDBJ database and are accessible via the BioProject ID: PRJDB17499.

### 2.6. Mapping of RNA-Seq reads

Raw sequencing data, produced by Illumina NovaSeq 6,000, were subjected to quality control, read filtering, read pruning, and adapter trimming using fastp ver 0.12.4 [13] with the default parameters. After filtering, the “clean reads” were mapped to the *Arabidopsis thaliana* TAIR10 genome using STAR ver. 2.7.10a [14] within the program RSEM ver. 1.3.1 [15], with the default parameters of RSEM selected. Multimapped reads were accepted unless the number of matched sites was more than 20. Mapped reads were quantified using RSEM. For differential expression analysis, genes with low counts (total counts across samples <=6) were filtered out, and raw counts of the remaining genes were normalized by the Trimmed-Mean of M values (TMM) method using the edgeR [16] option in the TCC package of R [17], with the following parameters set: iteration = 1, FDR = 0.1, and floorPDEG = 0.05.

### 2.7. Identification of differentially expressed genes (DEGs) using maSigPro

The R package maSigPro ver 1.66.0 [18] was used to identify DEGs in the timecourse RNA-seq data. The default parameters were adopted, with the exception of an *R^2^* threshold of ≥0.7. The identified DEGs were grouped into clusters according to expression patterns using maSigPro. To clarify the molecular and physiological responses to high concentrations of ammonium sulfate, functional enrichment analysis of the genes belonging to each cluster was conducted using the Metascape platform [19].

### 2.8. q-PCR analysis of gene expression in dde2-2 mutant treated with ammonium sulfate

The 21-day-old WT and *dde2-2* mutant seedlings were transferred onto the media containing 5 mM or 37.5 mM (NH_4_)_2_SO_4_ for 3 h, and leaves were harvested for RNA extraction. cDNAs were synthesized using PrimeScript RT Kit with gDNA Eraser (Perfect Real Time; Takara) following the protocol of the manufacturer. The relative expression of genes in leaves was determined by reverse transcription quantitative PCR performed on an Applied Biosystems StepOne Real-Time PCR System with KAPA SYBR® FAST qPCR Master Mix (2X) Kit (Kapa Biosystems) according to the protocol of the manufacturer. Primers used in the q-PCR analysis are listed in Table S1, and the expression data were normalized to *ACTIN2*.

## 3. Results

### 3.1. High ammonium sulfate treatment rapidly alters ammonium concentrations and the transcriptome in leaves

To check whether root-supplied ammonium led to leaf ammonium accumulation in a concentration-dependent manner, we first analyzed the time-course of ammonium concentrations in true leaves of plants treated with 5 mM (control treatment) or 12.5 mM and 37.5 mM (NH_4_)_2_SO_4_ (high ammonium sulfate treatments) (Fig. 1A and S1). High ammonium sulfate treatments increased ammonium concentrations in the leaves compared with the control treatment. In particular, 37.5 mM (NH_4_)_2_SO_4_ treatment rapidly increased the ammonium concentration, reaching approximately 3, 6, and 9 µmol g^−1^ at 3, 6, and 24 h after the treatment, respectively. After 24 h of the above treatments, and plants were transferred again to fresh 5 mM (NH_4_)_2_SO_4_ medium and grown for another 24 h. This treatment reduced leaf ammonium concentrations in 12.5 mM and 37.5 mM (NH_4_)_2_SO_4_-treated plants to similar levels of 5 mM control plants. These results indicate that the experimental method is suitable for investigating rapid plant responses to exogenous ammonium sulfate concentrations.

**Figure 1.**
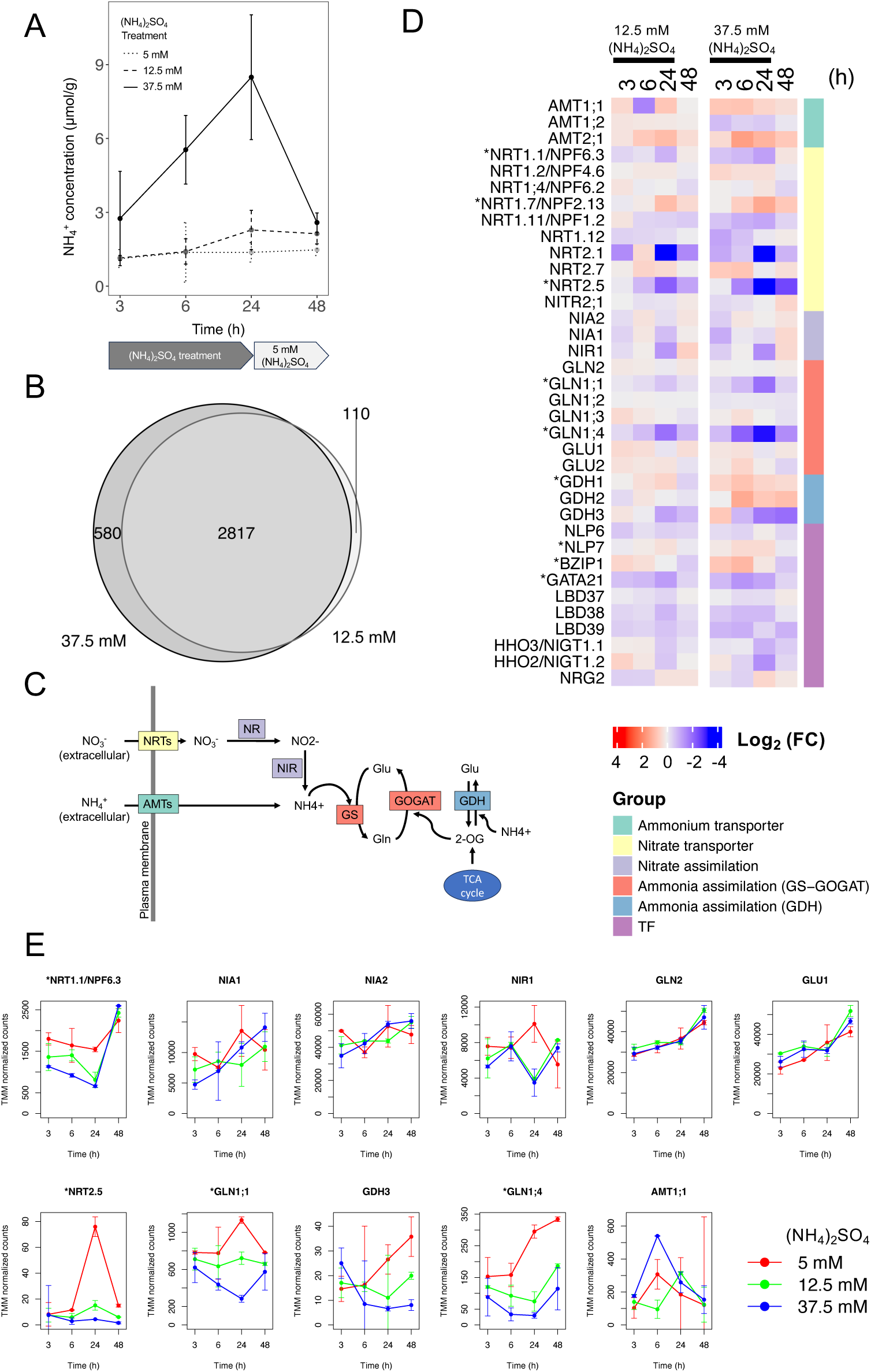
Elevated leaf ammonium concentrations and transcriptional changes in genes related to nitrate uptake and metabolism after high ammonium sulfate treatment. (A) Mean ammonium concentration (n = 2) of leaves during ammonium sulfate treatment (3 h, 6 h, and 24 h) and after being transferred back to cultivation medium with 5 mM (NH_4_)_2_SO_4_ (48 h). (B) Venn diagram showing the numbers of shared and unique DEGs identified in each pair of datasets (5 mM vs 12.5 mM or 37.5 mM (NH_4_)_2_SO_4_). (C) Nitrogen uptake and assimilation reaction pathway. NRT: nitrate transporter, AMT: ammonium transporter, NR: nitrate reductase, NIR: nitrite reductase, GS: glutamine synthetase, GOGAT: glutamate synthase, GDH: glutamate dehydrogenase, Gln: glutamine, Glu: glutamine, 2-OG: 2-oxoglutarate. The color coding of enzymes corresponds to the functional classification used in Fig. 1D. (D) Heatmaps of log2 fold change in the expression of the representative genes related to nitrogen metabolism for two different treatments (12.5 mM and 37.5 mM ammonium sulfate) at four time points. Log2 fold change was calculated based on normalized counts compared with the control samples (5 mM (NH_4_)_2_SO_4_). Color coding on the right side of the heatmap represents the functional classification of the genes. (E) Mean expression levels (n = 2) of nitrate inducible genes (*NRT1.1/NPF6.3*, *NIA1*, *NIA2*, and *NIR1*), N-starvation genes (*NRT2.5*, *GLN1;1*, *GLN1;4*, and *GDH3*, and *AMT1;1*), and ammonium assimilation genes (*GLN2* and *GLU1*). DEGs under the 37.5 mM treatment are marked with an asterisk (D and E).

To reveal transcriptomic alterations during ammonium sulfate treatment and post-treatment recovery, mRNAs were isolated from leaves and used for RNA-seq analyses. The maSigPro program identified 2,927 and 3,397 differentially expressed genes (DEGs) following the 12.5 and 37.5 mM (NH_4_)_2_SO_4_ treatments, respectively. Although most DEGs overlapped, 110 and 580 unique DEGs were identified in 12.5 mM and 37.5 mM (NH_4_)_2_SO_4_ treatments, respectively (Fig. 1B). In total, 3,507 DEGs identified in both treatments are listed in Table S2 along with their expression levels.

### 3.2. Nitrate-inducible genes and N starvation-inducible genes were downregulated during high ammonium sulfate treatment

Next, we examined the transcriptional changes in N transporters and assimilation genes during the treatments (Fig. 1C–E). The nitrate-inducible nitrate transporter genes *NRT1.1/NPF6.3* and *NRT2.1* [20] were downregulated by high ammonium sulfate treatment (Fig. 1D and E). However, little difference was observed in the expression of the nitrate-inducible nitrate reductase genes *NIA1* and *NIA2* among the treatments. The nitrite reductase gene *NIR1*, which is also nitrate-inducible, was transiently but strongly downregulated at 24 h after the treatments. Moreover, most of the known nitrate-inducible genes were either downregulated or unresponsive to high concentrations of ammonium sulfate (Table S3). These results suggest that high ammonium sulfate treatments generally cause attenuation of the nitrate-dependent induction of gene expression, although the transcription-activating nitrate sensor gene *NLP7* [21] was upregulated by 37.5 mM (NH_4_)_2_SO_4_ treatment (Fig. 1D).

Regarding ammonium assimilation genes, expression of the plastidic glutamine synthetase gene *GLN2* and ferredoxin-dependent glutamate synthase gene *GLU1* changed only slightly in response to different concentrations of ammonium sulfate (Fig. 1D and E). In contrast, high ammonium sulfate treatment significantly induced the expression of the glutamate dehydrogenase gene *GDH1*, suggesting increased assimilation of excessive ammonium via the GDH pathway.

Further detailed analysis of N-relevant genes found that some typical N starvation-inducible genes, such as the high-affinity nitrate transporter gene *NRT2.5* and ammonium assimilation genes *GLN1;1*, *GLN1;4*, and *GDH3* [22], were downregulated by high ammonium sulfate treatments (Fig. 1D and E). The N starvation-inducible ammonium transporter gene *AMT1;1* was exceptionally upregulated. Two groups of N-inducible transcription factors (TFs), LBD37/38/39 and NIGT1.1/1.2, act as repressors of N starvation-inducible genes in shoots [23,22]. These repressor genes were downregulated by 37.5 mM (NH_4_)_2_SO_4_ treatment (Fig. 1D). Collectively, these results show that under high concentrations of ammonium sulfate, N starvation-inducible genes could be down-regulated by a pathway distinct from LBD37/38/39 and NIGT1.1/1.2.

### 3.3. Clustering of DEGs identified in response to high ammonium sulfate treatments

The DEGs in the 12.5 mM and 37.5 mM (NH_4_)_2_SO_4_ treatments were classified into nine clusters by the maSigPro program based on their expression patterns (Fig. 2; Fig. S2 and S3). Information on the DEGs belonging to each cluster is summarized in Table S2. In these clusters, most of the changes in 12.5 mM (NH_4_)_2_SO_4_ treatments were positioned between the 5 and 37.5 mM treatments, indicating that the expression of those DEGs changed more as the (NH_4_)_2_SO_4_ concentration increased. Because most of the DEGs overlapped between the 12.5 mM and 37.5 mM treatments (Fig. 1B) and the clustering patterns were also similar between them (Fig. S2 and S3), we focused on the results of 37.5 mM (NH_4_)_2_SO_4_ treatment for further analysis.

**Figure 2.**
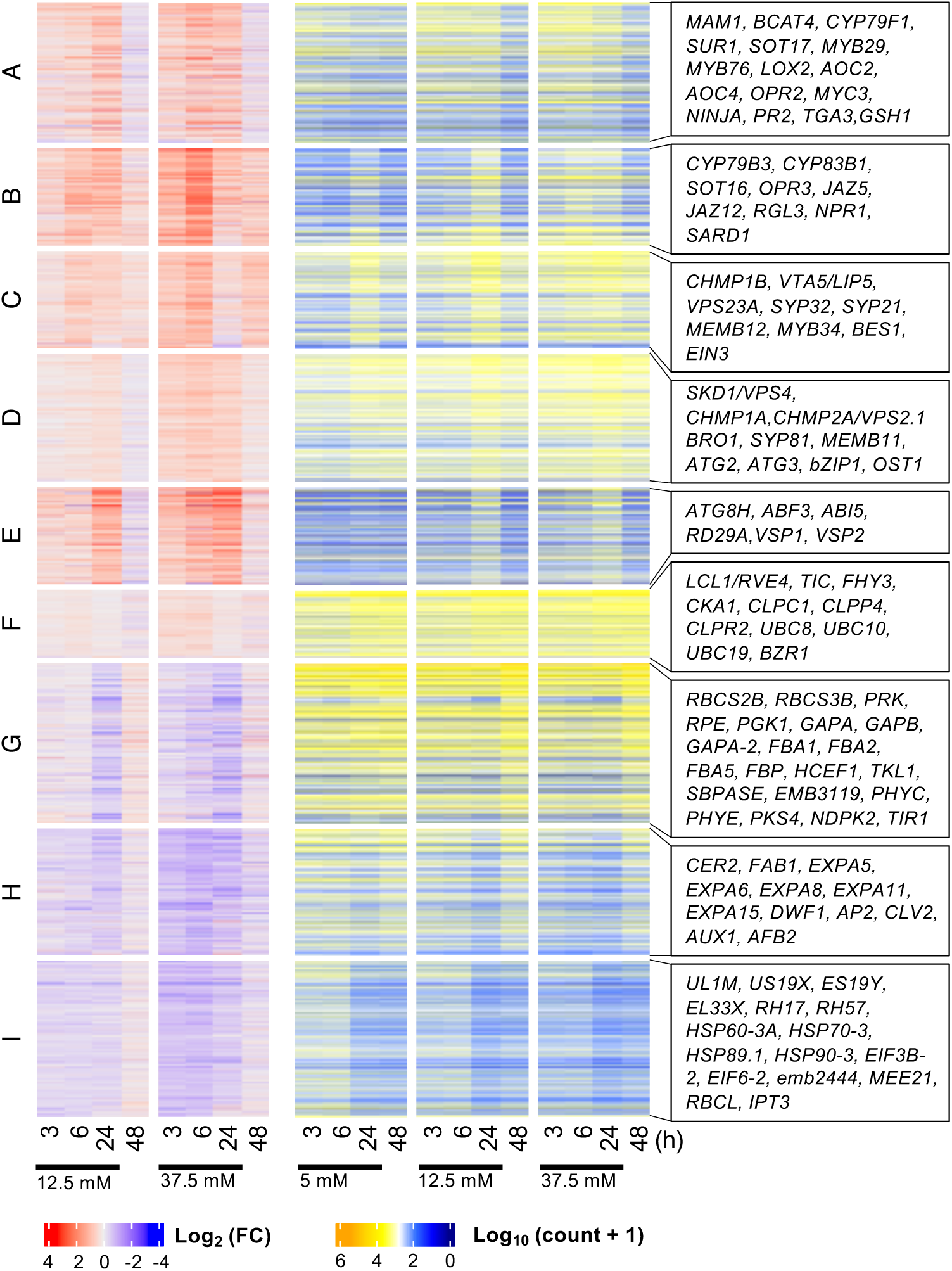
Transcriptional dynamics of identified DEGs in response to high (NH_4_)_2_SO_4_ treatment. The 3,397 DEGs identified under 37.5 mM ammonium sulfate treatment were divided into 9 clusters based on their temporal expression patterns using the maSigPro R package. The left and right heatmaps represent the change in expression (log2 fold change compared to the 5 mM (NH_4_)_2_SO_4_ condition) and expression level (log10 normalized counts) of the DEGs during ammonium sulfate treatment (3 h, 6 h, and 24 h) and post-treatment recovery (48 h), respectively. Representative genes for the functions enriched in each cluster are shown on the right.

The genes in clusters A–F were generally upregulated in the 37.5 mM (NH_4_)_2_SO_4_ treatment compared with the control, whereas those in clusters G–I were generally downregulated (Fig. 2 and Fig. S2). Clusters A and B included the genes that were most upregulated in the early stage of treatment (3 or 6 h), and clusters C–E included those that were upregulated in the late stage of treatment (24 h). In cluster F, gene expression at 3 h after the treatment was larger in 37.5 mM treatment than in the control, but thereafter it became similar between them. The expression of genes in clusters G and H was repressed during treatment. After the plants were returned to 5 mM medium, repression was lifted. The genes in cluster I remained repressed by 37.5 mM treatment even after the plants were returned to 5 mM medium.

### 3.4. JA- and SA-related defense response genes were upregulated in the early stages of high ammonium sulfate treatment

To determine the temporal physiological changes in response to 37.5 mM (NH_4_)_2_SO_4_ treatment, enrichment analyses of genes in each cluster were performed using Metascape [19] (Table S4 to S12).

The early upregulated genes in clusters A and B were overrepresented by genes related to the defense response to biotic stress (insect, bacteria, and fungi) (Table S4 and Table S5). The terms “S-glycoside biosynthetic process” and “defense response to other organism” were most significantly enriched in these cluster. The genes involved in the glucosinolate (GSL) biosynthetic process (e.g., *BCAT4*, *MAM1*, *CYP79F1*, *CYP79B3*, *CYP83B1*, *SUR1*, *SOT16* and *SOT17*) [24] and JA-responsive genes (such as the JA biosynthesis genes *LOX2*, *AOC2*, *AOC4*, *OPR2*, and *OPR3*, as well as the regulator genes *MYC3*, *NINJA*, *JAZ5*, *JAZ12*, and *RGL3*) [25] were included in these clusters. GSLs are a group of secondary metabolites that contribute to self-defense against insect herbivores in the plant order Brassicales [26]. In addition, genes related to salicylic acid (SA)-mediated immune responses (*PR2*, *TGA3*, *NPR1*, and *SARD1*) were included in these clusters (Fig. 2). JA mainly mediates the responses to wounding caused by insects and infection by necrotrophic microbes, whereas SA enhances resistance against biotrophic pathogens [27]. Given the widely accepted antagonistic interaction between JA and SA signaling, the dual upregulation of JA- and SA-related genes is noteworthy. One candidate for causing such an irregular response is NO, which is an endogenous signaling molecule crucial for plant responses to different stresses [28]. Previous studies have reported that NO induces the expression of many genes involved in oxidative stress and defense responses, including JA biosynthesis genes [29,30], SA-responsive genes [31], and genes involved in glutathione (GSH) metabolism [31,32], which correspond to the genes induced by high ammonium sulfate treatment in this study (Fig. 2 and Table S4). Furthermore, most NO-inducible genes [33] were upregulated in the early stage of high ammonium sulfate treatment (especially at 6 h) (Fig. S5). These results support the idea that endogenous NO might be involved in the transcriptional regulation in response to high concentrations of ammonium sulfate.

Since GSLs biosynthesis is regulated by JA [34], we investigated a potential role of JA signaling in ammonium tolerance and toxicity. We treated *dde2-2* mutant, impaired in JA biosynthesis [11], with ammonium sulfate. We found that the induction of *LOX2* and GSLs biosynthesis genes were attenuated in *dde2-2* mutant (Fig. 3). These results suggest that JA is essential for the induction of JA and GSLs synthesis genes by ammonium sulfate treatment.

**Figure 3.**
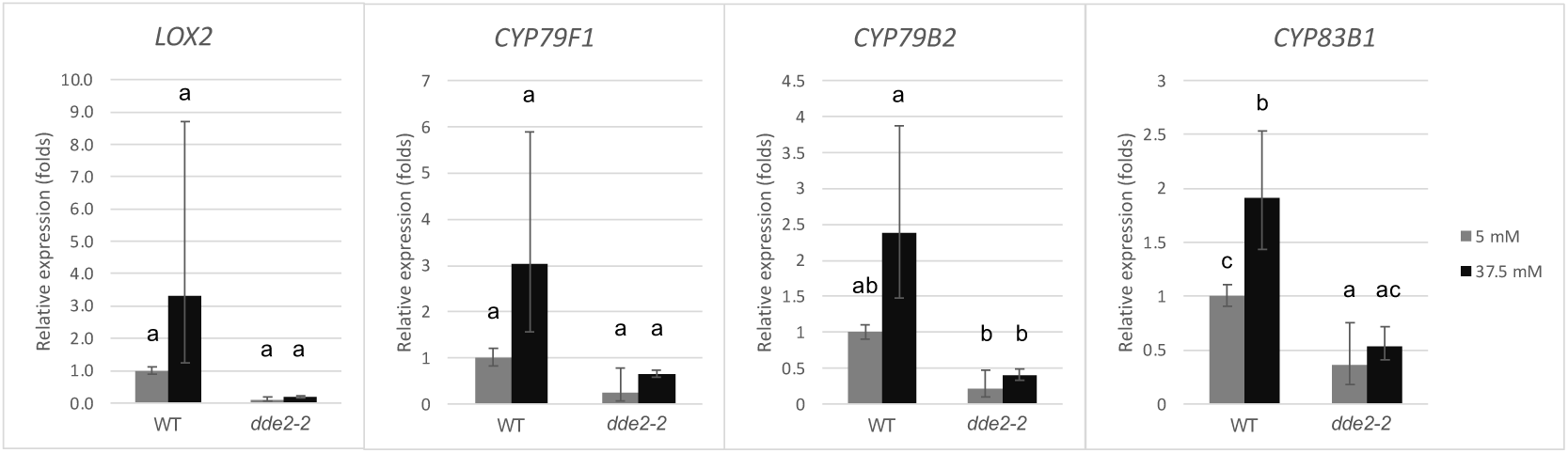
Gene expression levels of *LOX2* and GSLs biosynthetic genes by q-PCR in WT and *dde2−2* mutant in response to the (NH_4_)_2_SO_4_ treatment. Comparison of gene expression levels of JA biosynthetic gene *LOX2*, aliphatic GSLs biosynthetic gene *CYP79F1*, and indolic GSLs biosynthetic genes *CYP79B2* and *CYP83B1* between WT and *dde2-2* mutant with impaired JA synthesis, 3 hours after the treatment. Values were normalized to the expression of *ACTIN2* (mean ± SD; n = 2). Statistical analysis was conducted by Tukey-Kramer multiple comparisons test. Combinations with significant differences (*P* < 0.05) are labeled with different alphabetical symbols (a, b, and c).

### 3.5. Membrane trafficking-related genes and abscisic acid (ABA)/osmotic stress-related genes were upregulated during the late stage of high ammonium sulfate treatment

The late (24 h)-stage upregulated genes in clusters C and D were overrepresented by genes related to membrane trafficking, especially for ER–Golgi trafficking (e.g., *SYP81*, *SYP32*, *MEMB11*, *MEMB12*, *SYP21*), the vacuolar trafficking system including the late endosome/prevacuolar compartment/multi-vesicular body (e.g., *SKD1/VPS4*, *VTA1/LIP5*, *CHMP1A*, *CHMP1B*, *ALIX*, *CHMP2A, CHMP4, CHMP5, CHMP6, BRO1*, *VPS23A*), and autophagy (e.g., *ATG8, ATG2, ATG3, VPS15*, *ATI1*) (Table S6 and S7). Together with the upregulated ubiquitylation-related genes in cluster D (Table S7), these results suggest that the 24-hour high ammonium treatment stimulated an adaptation to a new environment through membrane remodeling and protein degradation. The metabolism of plasma membrane-localized nutrient transporters may be the targets of these upregulated genes.

The term “response to water deprivation” was most significantly enriched in cluster E (Table S8). The genes related to osmotic or salt stresses, such as abscisic acid (ABA)- independent TF *bZIP1*, and many ABA-related genes (e.g., *OST1, ABI5*, *ABF3*, and *RD29A*), were included in clusters D and E (Fig. 2, Table S7 and S8), indicating that osmotic stress responses were enhanced at the late stage.

### 3.6. Photosynthesis-related and growth-promoting genes were downregulated during high ammonium sulfate treatment

The DEGs in clusters G and H were downregulated during high ammonium sulfate treatment and recovered by 24 h after the end of treatment (Fig. 2 and Fig. S2). The term “photosynthesis” was most remarkably and significantly enriched in cluster G (Table S10). In addition, many terms related to photosynthesis (“reductive pentose-phosphate cycle,” “photosynthesis, light harvesting,” etc.) or chloroplast (“plastid organization,” “chloroplast localization,” etc.) were enriched in this cluster. This indicates that chloroplast activities were strongly inhibited during high ammonium sulfate treatment, but recovered after the treatment ended. Many genes encoding enzymes in the Calvin–Benson cycle were also found in cluster G, such as the Rubisco small subunit genes *RBCS*s, *RPE*, and *PRK* (Table S10). Furthermore, genes encoding phytochromes (*PHYC*, *PHYE*) and phytochrome signaling components (*PKS4*, *NDPK2*), which can fine-tune photomorphogenesis, were found in this cluster. In cluster H, the genes *CER2* and *FAB1* that contribute to fatty acid elongation, expansin genes (*EXPA*s) and *DWF1* that are involved in cell elongation, and *AP2* and *CLV2* that are related to development were included (Table S11).

Regarding the DEGs in cluster I, for which the expression levels recovered only slightly 24 h after the end of treatment, many genes encoding ribosomal proteins were enriched in this cluster. In addition, many heat shock protein genes (e.g., *HSP89.1*, *HSP60-3A*), translation initiation factor genes (*EIF*s), and embryo defective or arrest genes (e.g., *Emb2444*, *MEE21*) were included in this cluster (Table S12).

### 3.7. Genes associated with sulfate assimilation via PAPS are upregulated by high ammonium sulfate treatments

Several genes involved in sulfur assimilation had different expression patterns between 12.5 mM and 37.5 mM (NH_4_)_2_SO_4_ treatments (Fig. 4A and B). Although *SULTR1;2*, *APR* genes, and *SERAT3;2* were upregulated by the 12.5 mM treatment compared with the control, these genes were downregulated by the 37.5 mM treatment, but were then upregulated at 24 h after plants were returned to 5 mM (NH_4_)_2_SO_4_ medium. Meanwhile, the expression of *APK1* and *APK2* was significantly increased in both 12.5 and 37.5 mM treatments. APRs and APKs are involved in different sulfate assimilation pathways; the former mediates sulfite biosynthesis and the latter PAPS (3’- phosphoadenosine-5’-phosphosulfate) biosynthesis (Fig. 4A). Thus, the sulfate assimilation pathway via PAPS may be preferred during excess sulfate conditions (37.5 mM treatment).

**Figure 4.**
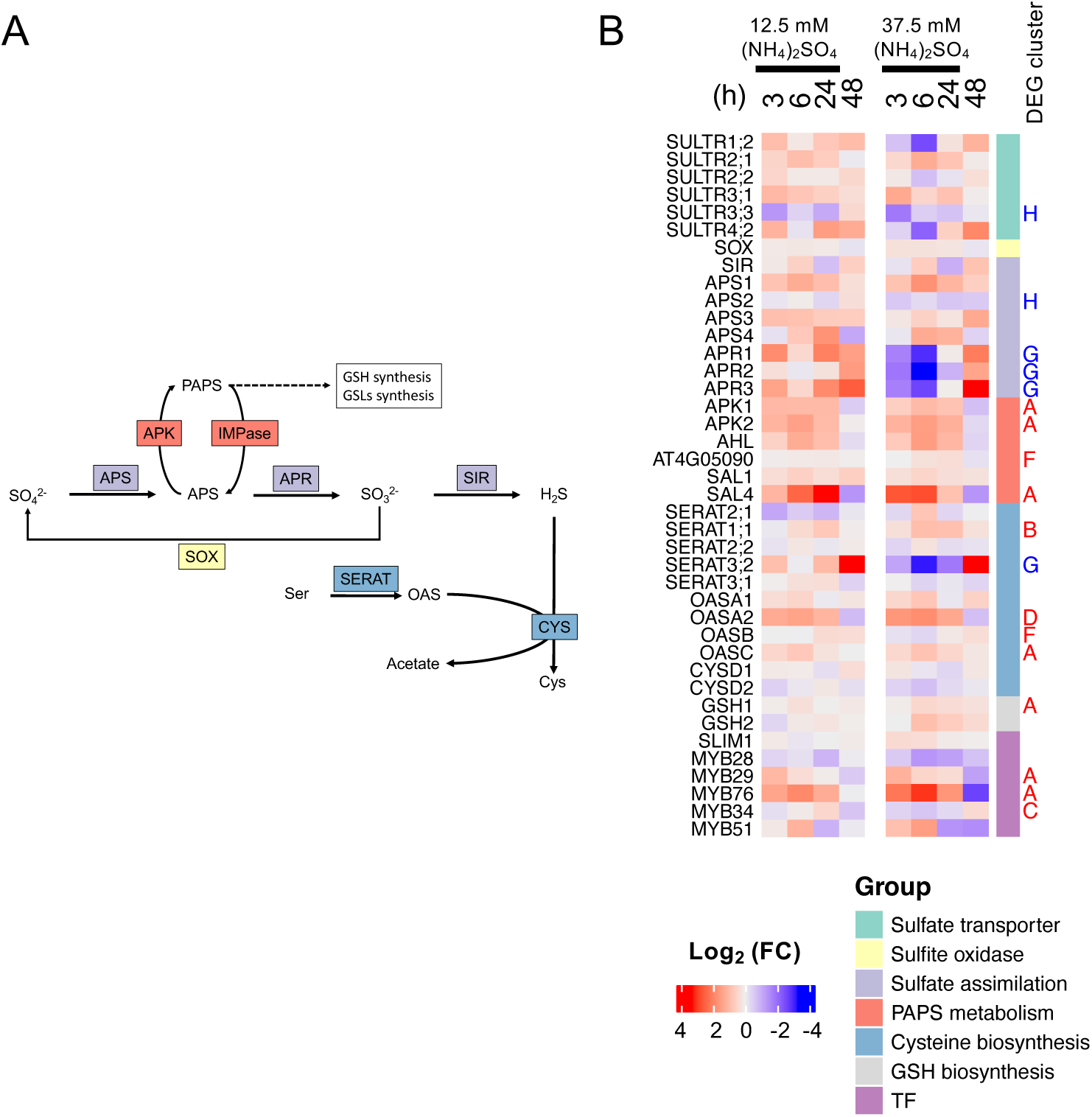
Upregulation of sulfate assimilation via PAPS in response to high-concentration ammonium sulfate treatment. A) Sulfate assimilation reaction pathway. APS (enzyme): ATP sulfurylase, APR: adenylyl-sulfate reductase, SIR: sulfite reductase, APK: APS kinase, IMPase: inositol monophosphatase, SOX: sulfite oxidase, SERAT: serine acetyltransferase, CYS: cysteine synthase, APS (compound): 5’-adenylylsulfate, PAPS: 3’-phosphoadenosine-5’- phosphosulfate, OAS: O-acetyl-L-serine, Ser: serine, Cys: cysteine, GSH: glutathione, GSLs: glucosinolates. B) Heatmaps of log2-fold change in the expression of representative genes related to sulfur metabolism. Explanations for the heatmaps are the same as those in Fig. 1D. For each DEG under 37.5 mM conditions, the DEG cluster it belongs to is denoted in red if it is upregulated and in blue if it is downregulated.

The expression of the TF gene *SLIM1*, for which the translational product upregulates the genes involved in sulfate uptake and assimilation in a sulfur-deficient environment [35], was not altered by high ammonium sulfate treatment (Fig. 4B). Among the GSL-associated R2R3-MYB TF genes, *MYB29* and *MYB76*, which induce aliphatic GSL biosynthesis [24,36], and *MYB34*, which induces indolic GSL biosynthesis [37], were detected as upregulated DEGs. In addition, the expression of the sulfate assimilation genes *APS1*, *APS3*, *APK1*, and *APK2* was induced by high ammonium sulfate treatments, which corresponded to a previous report that MYBs associated with GSLs can activate transcription of these genes [38]. APS and APK enzymes contribute to PAPS biosynthesis, and the high-energy sulfate in PAPS can be incorporated into GSLs (Fig. 4A). Thus, high ammonium sulfate treatments promote GSL biosynthesis through the simultaneous upregulation of GLS-associated *MYBs* and *APSs/APKs*.

### 3.8. Genes involved in the biosynthesis and signaling of phytohormones contributing to growth and development were downregulated during high ammonium sulfate treatment

The expression of genes related to auxin and cytokinin, phytohormones that promote growth and development, tended to be suppressed by high ammonium sulfate treatments (Fig. 5 and Fig. S4). Significant downregulation was observed for *AUX1*, which is responsible for auxin uptake into cells; *TIR1* and *AFB2*, which are involved in auxin signaling; and many auxin-responsive genes, including the auxin response factor family and the auxin/indole-3-acetic acid (*Aux*/*IAA*) family. Several genes involved in the biosynthesis of cytokinin, including *IPT3* and *CYP735A2*, were significantly down-regulated.

**Figure 5.**
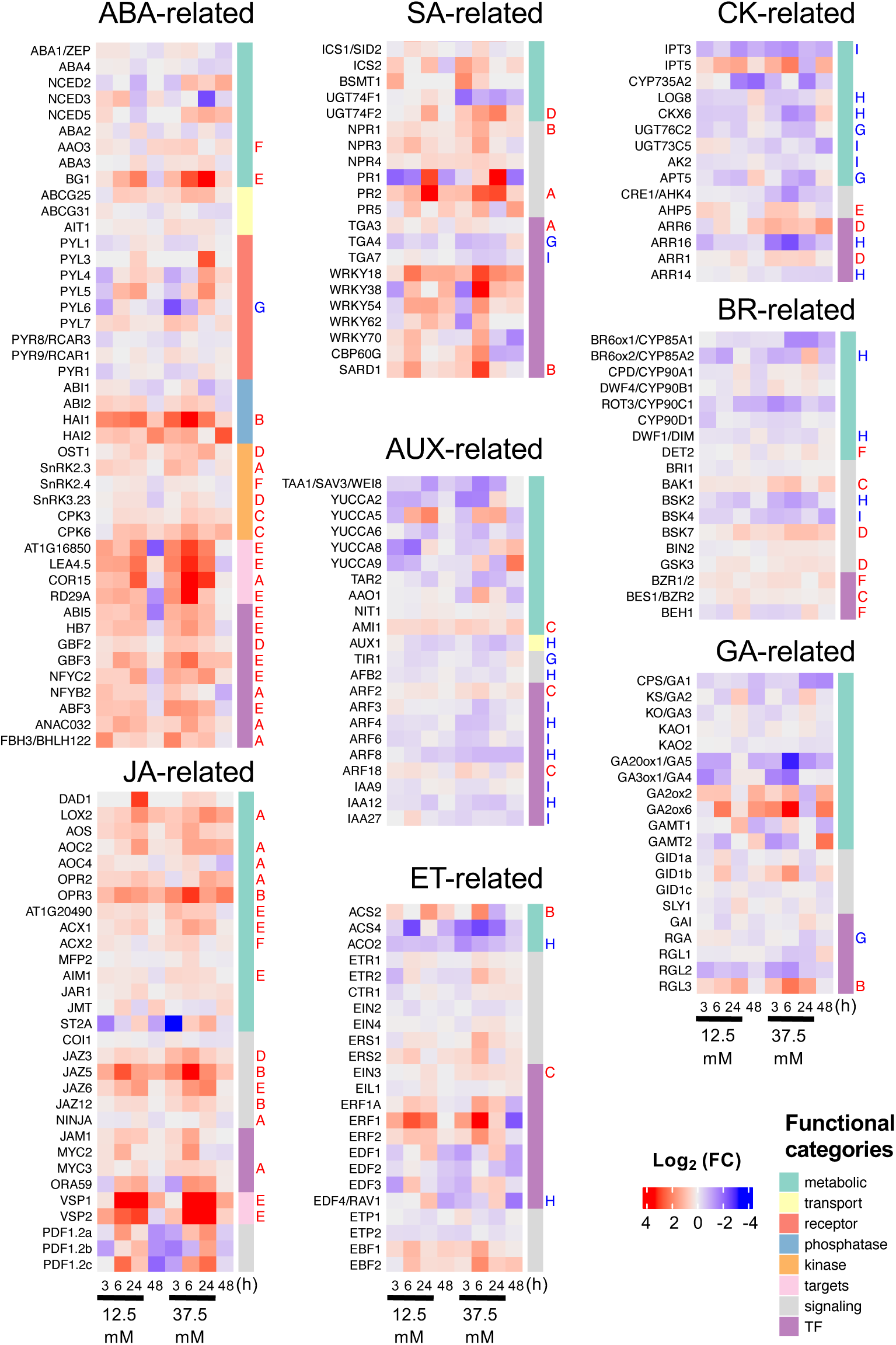
Phytohormone-related genes involved in the defense against abiotic and biotic stress were upregulated, whereas genes involved in growth and development were downregulated by high concentrations of ammonium sulfate. Log2 fold change in the expression of genes related to phytohormone metabolism and signaling. Explanations for the heatmaps are the same as those in Fig. 1D. For each DEG under 37.5 mM conditions, the DEG cluster it belongs to is denoted in red if it is upregulated and blue if it is downregulated DEG. Only selected representative genes or DEGs are shown. Heatmaps of all representative genes related to these hormones are shown in Fig. S4. ABA: Abscisic acid, JA: Jasmonic acid, SA: salicylic acid, AUX: auxin, ET: ethylene, CK: cytokinin, BR: brassinosteroid, GA: gibberellin.

For ethylene, brassinosteroids and gibberellins, no significant transcriptional changes were observed for most of the genes contributing to their biosynthesis and signaling. However, the *EIN3* gene, which encodes a TF initiating downstream transcriptional cascades for ethylene responses, and the *BZR1* and *BES1* genes, which encode TFs that are master regulators of brassinosteroid signaling, were significantly and most upregulated at 24 h after high ammonium sulfate treatment. The RGL3 gene, encoding a DELLA subfamily member that acts as a negative regulator of gibberellin signaling, was significantly upregulated. This suggested that the response to gibberellin, which acts on elongation growth, was suppressed.

Although post-translational regulations such as protein phosphorylation and ubiquitination are recognized as important for plant hormone signal transduction [39], we were also able to infer the roles of phytohormones underlying ammonium-induced growth inhibition from transcriptional changes.

## 4. Discussion

Nitrate and ammonium are the primary sources of N in soils. Although ammonium is a downstream metabolite produced during the process of nitrate assimilation, excess ammonium, unlike nitrate, causes toxic effects such as growth suppression and chlorosis. We identified more than 3,000 genes for which transcription levels were significantly altered by high concentrations of ammonium sulfate (Fig. 1B). These included genes involved in nutrition uptake, metabolism, membrane trafficking, and phytohormone responses, indicating that plants attempt to combat high concentrations of ammonium sulfate stress by regulating several processes.

Several studies have reported a possible interaction between ammonium and ABA in plants [40–42]. The upregulation of ABA-related genes observed in this study also showed that the ABA response was an important component of the response to ammonium sulfate (Fig. 5). Ammonium increases ABA accumulation in *Arabidopsis* roots [43] and facilitates ABA transport from root to shoot in *Ricinus* [40]. In our *Arabidopsis* leaf data, the expression of *NCED*s, which encode key enzymes in the de novo biosynthesis pathway of ABA, was not significantly altered; however, the expression of genes related to ABA signaling (*SnRK2*s and ABA-responsive TFs) and ABA-responsive genes (e.g., COR15 and RD29A) was upregulated (Fig. 5). *AAO3*, which is responsible for the final step of ABA synthesis, and *BG1*, which hydrolyzes and activates inactive ABA-glucose ester (ABA-GE), were upregulated in DEGs (Fig. 5). These results suggest that the ABA precursors and ABA synthesized in the roots could be transported to the leaves and activate ABA signaling in the leaves in response to high ammonium sulfate treatments.

Among the TFs involved in N metabolism expressed in leaves [44], *NLP7* and *bZIP1* were upregulated and *GATA21* was downregulated (Fig. 1D). NLP7 acts as a nitrate sensor to control the expression of nitrate-inducible genes [21]. Recently, it was found that this TF is transcriptionally upregulated by salt stress [45]. The expression of bZIP1, which is also induced by salt, especially in roots [46], can reverse many gene regulations, including N-inducible genes [47]. In addition, downregulated expression of the nitrate-inducible TF *GATA21* may be responsible for the downregulation of chloroplast and chlorophyll synthesis [48], and therefore might be related to chlorosis, a known symptom of ammonium toxicity. It is noteworthy that *ANAC032* was identified as an upregulated DEG gene in cluster A (Fig. 5). *ANAC032* expression is induced by phytohormones and abiotic/biotic stress (ABA, SA, methyl jasmonate, salt/osmotic stress, and infection with *Pseudomonas syringae* pv. *tomato* DC3000) [49,50]. ANAC032 reprograms carbon and nitrogen metabolism by increasing sugar and amino acid catabolism in plants during stress responses [51]. In ANAC032-overexpressing plants, photosynthetic activity was suppressed, resulting in the accumulation of reactive oxygen species (ROS) in chloroplasts and the induction of autophagy and ER stress response [51]. At least these TFs appeared to respond to stress induced by high concentrations of ammonium sulfate, regulating metabolism for survival, apparently resulting in outcomes such as growth inhibition and chlorosis, which are characteristic of ammonium toxicity.

We found that most NO-inducible genes [33] were upregulated in the early stage of high ammonium sulfate treatment (Fig. S5). In plants, NO can be biosynthesized via reductive (nitrite-dependent) and oxidative (arginine-dependent) pathways [52]. That is, NO can be generated from byproducts of both nitrate and ammonium assimilation. Furthermore, acidic conditions caused by an excess of ammonium may also activate non-enzymatic nitrite-dependent NO formation, as reported in the apoplast of barley grown in the presence of high concentrations of nitrate [53,54]. Therefore, it is quite plausible that endogenous NO was produced under the treatment conditions used in this study, where sufficient nitrate and excess ammonium were provided. Several reports have indicated the contribution of NO in tolerance to drought, salinity, temperature, and heavy metal stresses [28]. NO can counteract the effects of ROS produced by stresses [55] as well as alter the activity of various enzymes or TFs for stress tolerance through S-nitrosylation [56]. Interestingly, NLP7, a nitrate sensor and master regulator of primary nitrate response [21], was also found to be a target of S-nitrosylation [57]. Moreover, it has been reported that treatment with high doses (200 µM) of the NO donor SNAP abolishes the nuclear localization of NLP7 [58]. This might explain why the expression of NLP7 was upregulated, whereas the expression of nitrate-inducible genes was not promoted. Given that NO can react with GSH to produce S-nitrosothiols (SNO) and S-nitrosoglutathione (GSNO), which are NO donors for S-nitrosylation, the ammonium sulfate conditions in the present study, which promoted the expression of the GSH synthesis gene, may have promoted S-nitrosylation. It would be valuable to investigate the impact of NO and S-nitrosylation on ammonium toxicity using related mutants and NO donors/scavengers.

Collectively, our findings suggest a complex regulatory response to high ammonium sulfate concentrations, involving the modulation of genes related to nutrient uptake, metabolism, phytohormone signaling, and membrane trafficking. The upregulation of certain TFs associated with osmotic stress and NO-inducible genes could be an indication of the adaptive responses to the specific environmental conditions imposed by high ammonium sulfate treatment. Maintaining a balance between ammonium, nitrate, and sulfate appears to be crucial for fertilization with ammonium as the N source while reducing ammonium toxicity.

## Supplementary Materials

Figure S1. *A. thaliana* plants at 24 and 48 hours after the (NH_4_)_2_SO_4_ treatment.

Figure S2. Different time-series expression patterns of DEGs detected under 37.5 mM (NH_4_)_2_SO_4_ condition.

Figure S3. Different time-series expression patterns of DEGs detected under 12.5 mM (NH_4_)_2_SO_4_ condition.

Figure S4. Log2 fold change of the expressions of the genes related to phytohormone metabolism and signaling.

Figure S5. Log2 fold change of the expressions of the NO-inducible genes.

Table S1. Primer sequences for gene expression used in this study.

Table S2. List of the 3,507 DEGs, its cluster belongings and expression levels.

Table S3. Comparison of transcriptional change of genes regulated by nitrate or ammonium sulfate.

Table S4 – S12. List of grouped enriched terms as the result of enrichment analysis by Metascape for the DEGs in cluster A – I under 37.5 mM (NH_4_)_2_SO_4_ condition.

## Author Contributions

Conceptualization, H.I., Y.A., M.F., T.Hachiya and T.Hamada; Validation, H.I., Y.A., M.F., T.Hachiya and T.Hamada; Formal Analysis, H.I.; Investigation, H.I., Y.A., M.N., A.D., T.T., and T.Hachiya; Data Curation, H.I. and Y.A.; Writing – Original Draft Preparation, H.I.; Writing – Review & Editing, H.I., T.Hachiya and T.Hamada; Project Administration, M.F., T.Hachiya and T.Hamada; Funding Acquisition, T.Hachiya and T.Hamada

## Institutional Review Board Statement

Not applicable.

## Informed Consent Statement

Not applicable.

## Data Availability Statement

The data presented in this study are openly available in DDBJ database, BioProject ID: PRJDB17499.

## Acknowledgments

We would like to thank Dr. Hiroyuki Ashida (Shimane University) for technical support of ammonium determination.

